# Predictive Patterns of Antidepressant Response from Pre-Treatment Reward Processing using Functional MRI and Deep Learning: Key Results from the EMBARC Randomized Clinical Trial

**DOI:** 10.1101/2020.01.28.923821

**Authors:** Kevin P. Nguyen, Cherise Chin Fatt, Alex Treacher, Cooper Mellema, Crystal Cooper, Manish Jha, Benji Kurian, Maurizio Fava, Patrick J. McGrath, Myrna Weissman, Mary L. Phillips, Madhukar H. Trivedi, Albert Montillo

**Author notes:** **Address Correspondence to:** Albert Montillo, PhD, Director, Deep Learning for Precision Health Laboratory, Lyda Hill Department of Bioinformatics, University of Texas Southwestern Medical Center, Madhukar H. Trivedi, M.D. Professor of Psychiatry, Julie K. Hersh Chair for Depression Research and Clinical Care Betty Jo Hay Distinguished Chair in Mental Health, Director, Center for Depression Research and Clinical Care University of Texas Southwestern Medical Center. Drs. Montillo and Trivedi contributed equally as senior authors to this work.

## Abstract

*Importance:* The lack of antidepressant-specific biomarkers to inform treatment selection is a key obstacle in the treatment of Major Depressive Disorder. Quantitative measurements of reward processing neural activity with task-based functional magnetic resonance imaging (fMRI) may allow prediction of individual outcomes for specific antidepressants.

*Objective:* To build and validate predictors of individual outcomes for sertraline, bupropion, and placebo using reward processing task-based fMRI, clinical assessments, and deep learning.

*Design:* This is a secondary analysis of data from the Establishing Moderators and Biosignatures of Antidepressant Response in Clinical Care (EMBARC) study, a placebo-controlled, double-blind randomized clinical trial. The study ran from July 2011 to December 2015 and this analysis was performed between December 2018 and July 2019.

*Setting:* The EMBARC study was conducted at 4 academic medical centers.

*Participants:* A random sample of 296 un-medicated participants meeting DSIM-IV criteria for depression was enrolled.

*Intervention:* Subjects were randomized to sertraline or placebo during phase 1. In phase 2, non-responders to placebo were treated with sertraline and non-responders to sertraline were treated with bupropion. Each phase lasted for 8 weeks.

*Main outcomes and measures:* The primary outcome was the change in the 17-item Hamilton Rating Scale for Depression (ΔHAMD) after 8 weeks of treatment. Task-related brain activation measures, computed from pre-treatment reward processing task fMRI, and clinical measurements were used to train deep learning predictive models for each treatment group.

*Results:* The current analysis includes 222 participants with complete imaging, clinical, and treatment outcome data (146 female, mean age 16.4 ± 13.4 years). The predictive model for sertraline, trained on 106 participants, achieved an *R*^2^ of 35% (95% CI 20-50%, *p* < 10^−3^) in predicting ΔHAMD and a number-needed-to-treat (NNT) of 4.31 in predicting remission. The placebo model, trained on 116 participants, achieved an *R*^2^ of 23% (95% confidence interval of 11-37%, *p* < 10^−3^) and an NNT of 2.78. The bupropion model achieved an *R*^2^ of 37% (95% CI 12-61%, *p* < 10^−3^) and an NNT of 2.35. Reward processing activity in regions such as the medial frontal cortex, insula, and thalamus were important predictors of sertraline outcome, while the anterior cingulate cortex, striatum, insula, and thalamus were important regions for predicting bupropion outcome. These are consistent with previously reported findings, while regions not previously implicated such as the temporal pole and paracentral lobule were also found to be predictive.

*Conclusions and relevance:* These findings demonstrate the utility of reward processing measurements and deep learning in predicting individual antidepressant outcomes with high accuracy. They also present potential composite biomarkers for these treatments based on neuroimaging and clinical features.

*Trial registration:* ClinicalTrials.gov identifier: <u>NCT01407094</u>

**KEY POINTS:** *Question:* Can pre-treatment fMRI measurements of reward processing, in combination with multi-dimensional clinical assessments, be used to form personalized predictions of antidepressant outcomes?

*Findings:* Deep learning predictive models trained on pre-treatment reward processing task-based neuroimaging and clinical data from a randomized clinical trial were able to explain up to 37% of the variance in individual treatment outcomes and predict remission with NNT of 2-4. Specific clinical variables and brain regions with reward processing activity important for the model predictions were identified, which may form composite biomarkers of antidepressant response.

*Meaning:* Quantitative measures of reward processing, as identified through deep learning, may move us closer to a precision medicine approach that enables clinicians to select the appropriate antidepressant for each patient with greater certainty.

## INTRODUCTION

The discovery of biomarkers of antidepressant response is crucial to achieving personalized treatment planning in Major Depressive Disorder (MDD). Currently, remission rates for individual antidepressants are typically below 40%^1^, and about 33% of patients require over 3-4 drug trials before achieving remission^2^. However, biomarkers and predictive tools that could identify upfront the optimal antidepressant for each patient would reduce the need for multiple drug trials, expedite remission, and enable a precision medicine approach to MDD treatment.

Noninvasive measurements of individual brain activity patterns, such as those provided by functional magnetic resonance imaging (fMRI), have shown promise for this providing pre-treatment markers of treatment outcome. For example, analysis of resting-state fMRI has correlated default mode network and hippocampal connectivity with clinical severity change with sertraline treatment^3^. Additionally, fMRI measurements of the serotonergic emotional regulation circuit have revealed associations with treatment outcome. Greater anterior cingulate cortex activation during emotional regulation has been associated with improved response to venlafaxine and fluoxetine^4, 5^. Amygdala activation during exposure to emotional stimuli has similarly been connected to non-specific treatment outcome^6^. The dopaminergic reward processing circuity has been less studied for treatment outcome prediction, though one recent study has identified abnormal reward learning activity in the ventral striatum as a predictor of better response to sertraline over placebo^7^. There has also been little previous work on bupropion response biomarkers. One fMRI study of bupropion has been conducted at this time^8^, which examined emotional regulation rather than the dopaminergic reward processing circuitry primarily modulated by this drug^9^.

While this previous work highlights the potential of neuroimaging to predict treatment response, they are limited by the statistically complex nature of fMRI data. Deep learning models may be more apt to discover the complex, nonlinear association between fMRI measurements and treatment outcome. Also, the capability of deep learning models to scale to large numbers of input features, while automatically learning the most informative ones, is exploited to integrate imaging features with pre-treatment clinical measurements.

This secondary analysis of the EMBARC study explored whether pre-treatment reward task-based fMRI could be used to predict treatment-specific outcome using a data-driven, depp learning approach. Separate models were created for each of the 3 treatment arms (sertraline, placebo, and bupropion). The predictive performance of each model is reported, along with the most predictive features for each treatment. These predictive features may serve as candidate biomarkers of treatment outcome, and they suggest specific aspects of reward processing and clinical phenotype that are most pertinent for personalized treatment planning.

## METHODS

### Participants

This work is a secondary analysis of the Establishing Moderators and Biosignatures of Antidepressant Response in Clinical Care (EMBARC) study. The full study design has been previously reported^9^. A total of 296 participants with MDD were enrolled with written informed consent and institutional review board approval across 4 study sites. Inclusion criteria included early onset (before age 3) and chronic (episode duration > 2 years) or recurrent (2+ episodes) disease. Further details on inclusion/exclusion criteria (Appendix I), demographics (Table S1), and a CONSORT flow diagram (Figure S1) are included in the Supplement.

### Treatment Protocol and Outcomes

The treatment period included two 8-week phases. In Phase 1, participants were randomized under double-blind conditions into sertraline or placebo treatment arms. Randomization was stratified by study site, baseline depression severity, and disease duration. At week 8, sertraline-treated participants not meeting response criteria (Clinical Global Improvement score less than “much improved”) were crossed over to bupropion treatment in Phase 2. Clinical severity was tracked using the 17-item Hamilton Rating Scale for Depression (HAMD), and the primary outcome in this analysis is the change in HAMD (ΔHAMD) over the 8-week treatment phase (week 8 minus baseline for sertraline and placebo, week 16 minus week 8 for bupropion). Additional binary outcomes were defined using standard clinical criteria, including *response* (decrease in HAMD ≥ 50% from pre-treatment) and *remission* (week 8 HAMD ≤ 7). Mean ΔHAMD, response rates, and remission rates for each treatment arm are presented in Table S1 of the Supplement.

### MRI Acquisition

Functional and structural MRI scanner and sequence information for each study site can be found in Table S2 of the Supplement. Differences among the sites were controlled by the fMRI processing pipeline described below, which computes normalized brain activation measurements.

Reward task-based fMRI was acquired for 8 minutes during a block-design number-guessing task which probes reward processing neural circuitry known to be altered in MDD (Supplement, Appendix I and Figure S2)^10, 11^. Briefly, the task comprises 24 trials with random outcomes, 12 of which are “possible win” trials (50% chance of reward) and 12 of which are “possible loss” trials (50% chance of punishment). Each trial consists of an anticipation phase (subject is informed about the trial type), response phase (subject guesses whether the upcoming number is > or < 5), outcome phase (the number is revealed, and the resulting reward or punishment is displayed), and an inter-trial baseline phase.

Images with focal signal loss and clipped field-of-view were removed, and final inclusion criteria for this analysis were completion of 8 weeks of a given treatment. This yielded 106 participants with fMRI and sMRI for sertraline, 116 for placebo, and 37 for bupropion.

### Data Augmentation

Data augmentation is commonly employed in deep learning to increase performance and the likelihood of learning true associations when data is limited. This technique applies randomly-generated geometric transformations to real images to simulate additional images. The approach employed here uses co-registrations between participant sMRI data to simulate variations in brain morphology, synthesizing new fMRI acquisitions from the original data^12^. Augmentation was applied 5 times for each original image, providing a total of 222 fMRI images for bupropion, 978 for sertraline, and 696 for placebo. Importantly, this augmented data was used only during model training and not during evaluation. Additional results in Table S5 in the Supplement demonstrate the performance benefit achieved with vs. without augmentation.

### MRI Preprocessing and Feature Extraction

Original and augmented fMRI were preprocessed using conventional steps, including skull-stripping, head motion correction, spatial normalization, and spatial smoothing (Supplement, Appendix I). To quantify relative brain activation during task conditions, 3 contrast maps were computed for each participant. These contrasts included 1) *anticipation*, representing the initial phase of each trial in which the subject is informed whether the trial is potentially rewarding or punishing; 2) *reward expectation*, representing the contrast between anticipation of rewarding vs. punishing trials; and 3) *prediction error*, representing the contrast between the outcome phase for correct vs. incorrect guesses. From each contrast map, 200 mean regional values were extracted using a study-specific brain parcellation (Supplement, Appendix I), yielding 600 fMRI features for each participant.

### Acquisition of Clinical Measurements

In addition to imaging features, 95 pre-treatment clinical measures and demographic features were also included as predictor inputs. Demographics consisted of race, ethnicity, age, education, biological sex, and marital status. Clinical measures included total scores and sub-scores for several subject-reported forms and clinician-administered assessments (Supplement, Table S3). These included measurements of pre-treatment depression severity, childhood trauma, mania, anger attacks, suicide risk, anhedonia severity, anxiety severity, episode duration, nicotine dependence, and personality traits. Participant and family psychiatric history items were also included.

### Deep Learning Model Training and Evaluation

Feed-forward neural networks were constructed to take parcellated contrast map values and clinical features as inputs and return the predicted ΔHAMD. A separate model was trained for each treatment: sertraline, bupropion, and placebo. Models were evaluated using nested cross-validation, which provides an unbiased estimate of real-world predictive performance^13^, with 3 outer and 5 inner folds. Full technical details of the model architecture, hyperparameter optimization, nested cross-validation, and implementation are available in the Deep Learning Model Training and Hyperparameter Optimization section and Figure S3 of the Supplement.

Accuracy metrics for predicting ΔHAMD included R^2^ (coefficient of determination) and root mean squared error (RMSE), computed between model-predicted and true ΔHAMD. The model ΔHAMD predictions were then thresholded to obtain predictions of the two binary outcomes, remission and response. Accuracy metrics for predicting remission and response included predictive value (PPV), and area under the receiver operating characteristic curve (AUROC), and number-needed-to-treat (NNT). Permutation testing was performed to test the statistical significance of each model’s performance, and the most important features learned by each model to predict treatment outcome were identified using permutation feature importance (Supplement, Appendix I)^14^. In the Results, the top 30 features are reported for each model as the subsequent features had negligible permutation feature importance.

## RESULTS

### Prediction of Sertraline Treatment Outcome

For the sertraline treatment arm (*n* = 106), the mean ΔHAMD was 7.89 ± 7.16, remission rate was 39%, and response rate was 54% (Supplement, Table S1). The sertraline predictive model achieved a substantial *R*^2^ of 35% (95% confidence interval of 20-50%) and RMSE of 5.75 in predicting ΔHAMD (**Table 1**, 1^st^ row). NNT was 4.31 and PPV was 62% for predicting remission, and NNT was 4.88 and PPV was 68% for predicting response. Permutation testing confirmed these results to be statistically significant with *p* < 10^−3^. Notably, the best performance was reached when trained on the combination of imaging and clinical/demographic features versus using imaging features (*R^2^* 2%) or clinical features (*R^2^* 2%) alone (Supplement, Tables S5 and S6).

**Table 1.** Outcome prediction performance for the three treatments investigated. Deep learning models were trained to predict 8-week ΔHAMD. Performance metrics for this target include the coefficient of determination (*R*^2^) and root mean squared error (RMSE). To obtain predictions of remission and response, which are binary variables, model outputs were thresholded post-hoc using the HAMD criteria for remission (HAMD ≤ 7 at week 8) and response (decrease in HAMD ≥ 50%). Performance metrics for remission and response are number-needed-to-treat (NNT), positive predictive value (PPV) and area under the receiver operating characteristic curve (AUROC). Statistical significance of these performance measurements over chance accuracy was measured using permutation testing, and the *p*-values are presented here.

| Treatment | Features used | Prediction target |  |  |  |  |  |  |  |
| --- | --- | --- | --- | --- | --- | --- | --- | --- | --- |
| | | ΔHAMD | | Remission | | | Response | | |
| | | $R^2$ | RMSE | NNT | PPV | AUROC | NNT | PPV | AUROC |
| Sertraline | Imaging, clinical | 35%<br>$p < 10^{-3}$ | 5.75<br>$p < 10^{-3}$ | 4.31<br>$p < 10^{-3}$ | 0.62<br>$p < 10^{-3}$ | 0.60<br>$p=0.017$ | 4.88<br>$p < 10^{-3}$ | 0.68<br>$p < 10^{-3}$ | 0.63<br>$p < 10^{-3}$ |
| Placebo | Imaging, clinical | 23%<br>$p < 10^{-3}$ | 6.06<br>$p < 10^{-3}$ | 2.78<br>$p < 10^{-3}$ | 0.69<br>$p < 10^{-3}$ | 0.61<br>$p=0.048$ | 3.19<br>$p < 10^{-3}$ | 0.67<br>$p < 10^{-3}$ | 0.64<br>$p < 10^{-3}$ |
| Bupropion | Imaging | 37%<br>$p < 10^{-3}$ | 4.36<br>$p < 10^{-3}$ | 2.35<br>$p=0.018$ | 0.75<br>$p=0.044$ | 0.71<br>$p=0.196$ | 3.24<br>$p=0.002$ | 0.71<br>$p=0.002$ | 0.62<br>$p=0.006$ |

Of the 30 features with the highest permutation feature importance, 24 were imaging features and 6 were clinical and demographic features (**Fig. 1a**), with the clinical features ranking most highly. Of the clinical features, higher pre-treatment symptomatic severity (17-item and 24-item HAMD total), body mass index (BMI), psychomotor retardation reported on the SCID^a^ assessment, and higher NEO^b^ Neuroticism score were predictive of <u>resistance</u> to treatment (no remission). Psychomotor agitation reported on the SCID assessment was predictive of <u>remission</u>.

**Figure 1.**
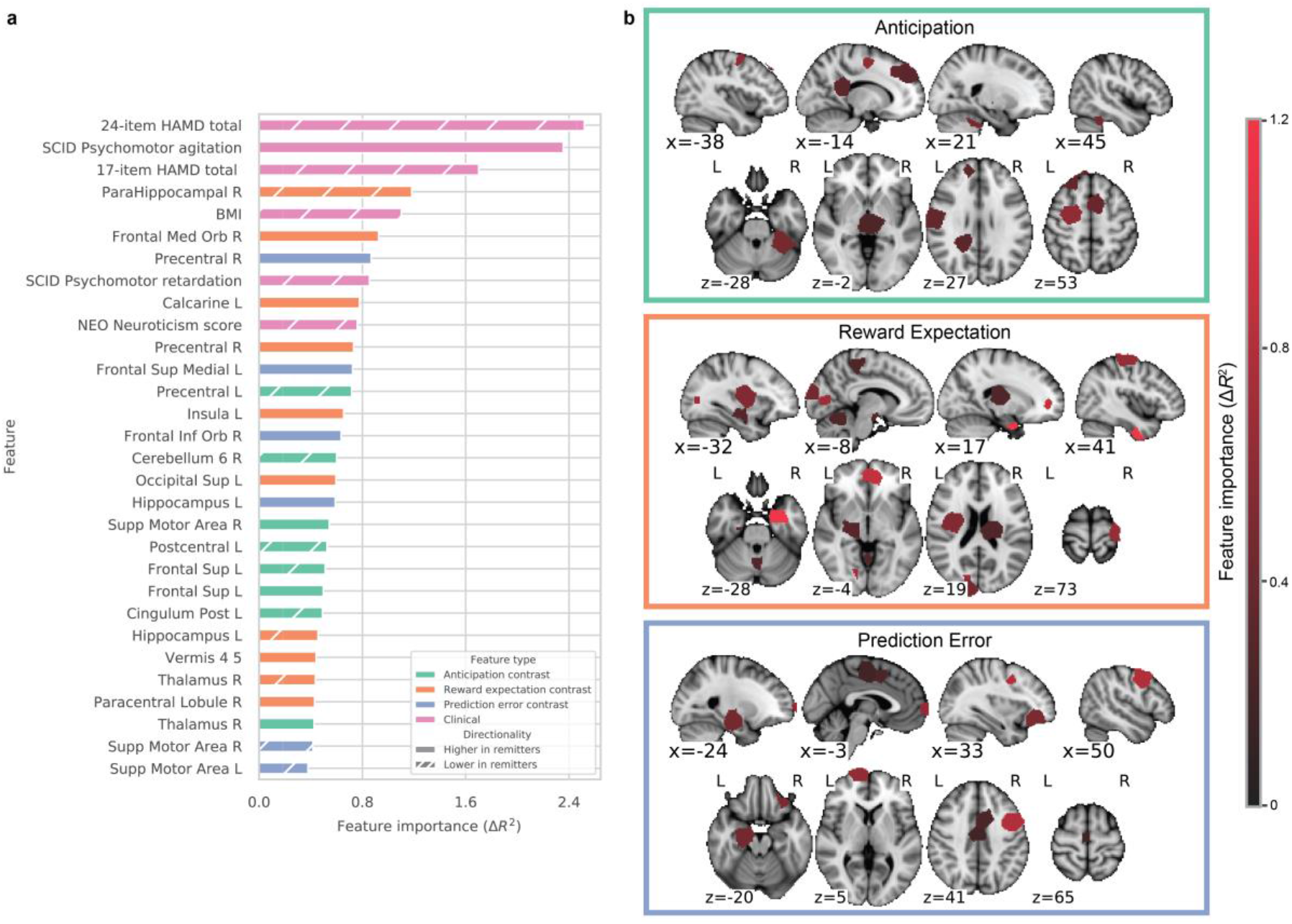
The thirty most important features learned by the sertraline outcome prediction model, as determined by permutation feature importance. **a)** Features are ranked by descending importance and colored by feature type. The directionality of each feature was computed by comparing the mean feature values of remitters and non-remitters; features with lower values in remitters have hashed bars. **b)** The imaging features are visualized as colored ROIs in the study-specific brain atlas and overlaid on the MNI brain template. Brighter red indicates greater feature importance.

Examining the imaging features (**Fig. 1b**), resistance was predicted by higher BOLD response to reward expectation in the parahippocampal gyrus, hippocampus, and thalamus; higher BOLD response to prediction error in the supplemental motor area; and higher BOLD response to anticipation in the precentral gyrus, anterior cerebellum, postcentral gyrus, and superior frontal gyrus. Remission was predicted by higher BOLD response to reward expectation in the medial orbitofrontal cortex, calcarine sulcus, precentral gyrus, insula, superior occipital lobe, cerebellar vermis, and paracentral lobule; higher BOLD response to prediction error in the precentral gyrus, medial superior frontal gyrus, inferior orbitofrontal cortex, and hippocampus; and higher BOLD response to anticipation in the supplemental motor area, superior frontal gyrus, and thalamus.

### Prediction of Placebo Treatment Outcome

In the placebo treatment arm (*n* = 116), the mean ΔHAMD was 6.70 ± 6.93, not significantly different from the mean ΔHAMD of the sertraline arm (*p* = 0.11). Remission and response rates were 33% and 35% respectively, lower than those of sertraline (Supplement, Table S1). The placebo model attained *R*^2^ of 23% (95% confidence interval of 11-37%) and RMSE of 6.06 for predicting ΔHAMD on held-out test data (**Table 1**, 2^nd^ row). NNT was 2.78 and PPV was 69% for predicting remission, and NNT was 3.19 and PPV was 67% for predicting response. This performance was statistically significant (*p* < 10^−3^) upon permutation testing. Like the sertraline model, the highest predictive performance was achieved using both imaging and clinical/demographic features versus imaging (*R^2^* 7%) or clinical features (*R^2^* 2) alone (Supplement, Tables S5, S6).

Compared to the sertraline model, the placebo model relied more heavily on clinical and demographic features. Of the 30 most important features learned by the model (**Fig. 2a**), 15 were imaging features and 15 were clinical and demographic features. Features predictive of <u>resistance</u> included: concurrent anxious distress and panic disorder (from SCID), greater anhedonia (MASQ^c^ anhedonic depression score and SHAPS^d^ ordinal total), number of comorbidities (SCQ total), Caucasian race, age, the presence of melancholic, atypical, or catatonic depression (*SCID current episode specifier*), number of children, current alcoholism, and psychomotor agitation. <u>Remission</u> was predicted by Asian race, longest period without dysphoria, the Concise Association Symptom Tracking (CAST) scale, and concurrent psychotic symptoms.

**Figure 2.**
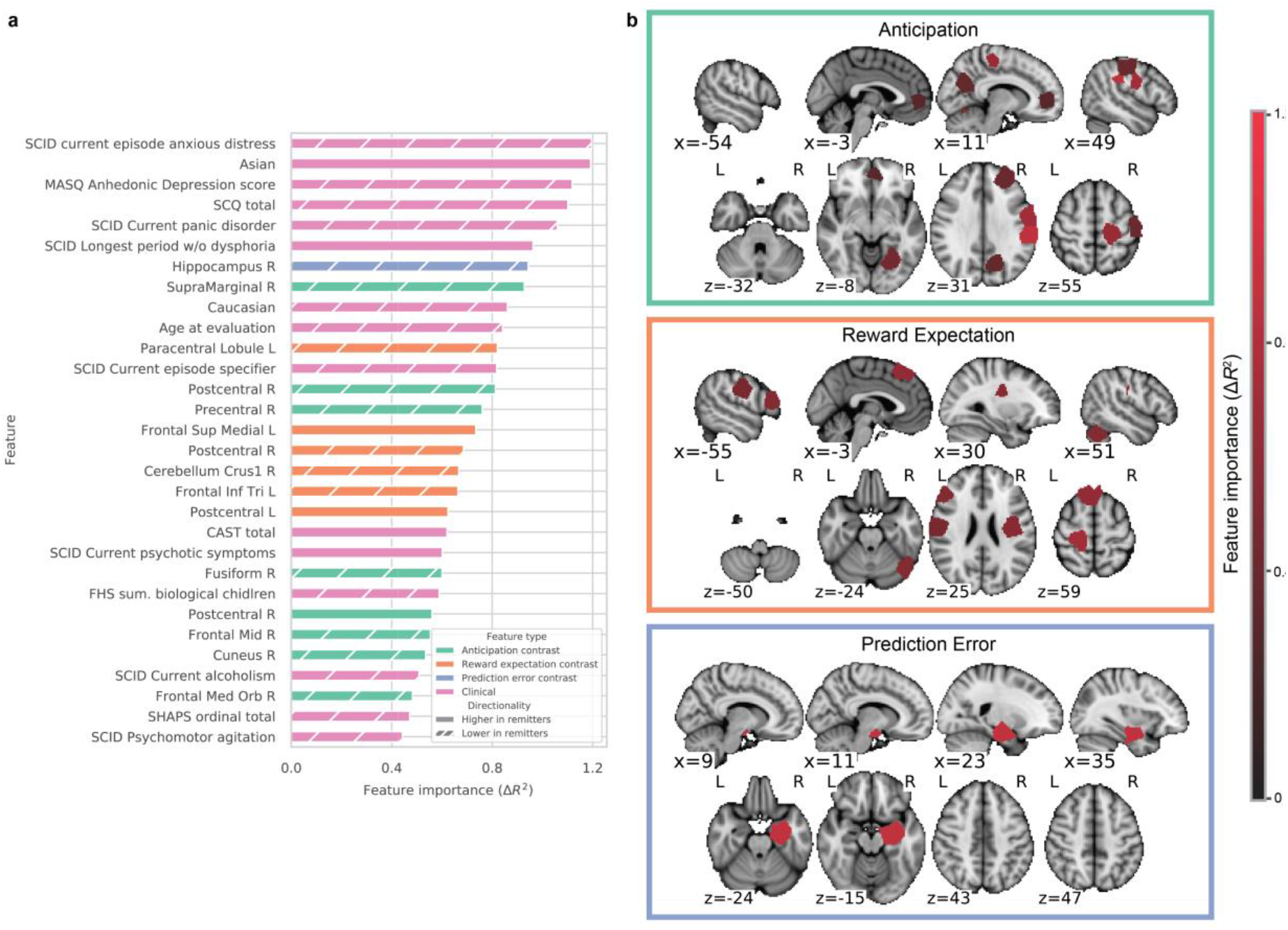
The thirty most important features learned by the placebo outcome prediction model, as determined by permutation feature importance. **a)** Features are ranked by descending importance and colored by feature type. The directionality of each feature was performed by comparing the mean feature values of remitters and non-remitters; features with lower values in remitters have hashed bars. **b)** The imaging features are visualized as colored ROIs in the study-specific brain atlas and overlaid on the MNI brain template. Brighter red indicates greater feature importance.

The imaging features were distinct from those learned by the sertraline model (**Fig. 2b**). Resistance was predicted by higher BOLD response to prediction error in the hippocampus; higher BOLD response to anticipation in regions near the central sulcus (supramarginal gyrus, postcentral gyrus, and precentral gyrus), the fusiform gyrus, middle frontal gyrus, cuneus, and medial orbitofrontal cortex; and higher BOLD response to reward expectation in the paracentral lobule, postcentral gyrus, triangular part of the inferior frontal gyrus, and cerebellar crus. Remission was predicted by higher BOLD response to anticipation in the postcentral gyrus and higher BOLD response to reward expectation in the medial superior frontal gyrus and postcentral gyrus.

### Prediction of Bupropion Treatment Outcome

At the end of Phase 1 of the study, subjects who failed sertraline were switched to bupropion for the Phase 2. Accordingly, ΔHAMD for bupropion was computed as the difference between the end and start points of Phase 2 (weeks 16 and 8). Mean ΔHAMD was 5.46 ± 5.57 remission rate was 32%, and response rate was 41% for bupropion-treated participants (**Table 1**). In contrast to the sertraline and placebo models, the bupropion model achieved the highest performance in predicting ΔHAMD using imaging features alone; the addition of clinical and demographic features was found to decrease performance (*R^2^* 25%, Supplement, Table S5). This imaging-only model achieved *R*^2^ of 37% (95% confidence interval of 12-61%) and RMSE of 4.36 in predicting ΔHAMD, with *p* < 10^−3^ upon permutation testing (**Table 1**, 3^rd^ row). NNT was 2.35 (*p* = 0.018) and PPV was 75% (*p* = 0.044) for predicting remission, and NNT was 3.24 (*p* = 0.002) and PPV was 71% (*p* = 0.002) for predicting response.

The important imaging features of the bupropion model (**Fig. 3**) were distinct from those learned by the sertraline and placebo models. Resistance was predicted by higher BOLD response to reward expectation in the frontal operculum and inferior parietal gyrus and higher BOLD response to prediction error in the lingual gyrus, postcentral gyrus, superior parietal gyrus, superior and middle frontal gyri, Rolandic operculum, supplemental motor area, rectus and thalamus. Remission was predicted by higher BOLD response to reward expectation in the posterior cingulate cortex, fusiform gyrus, middle occipital gyrus, and parahippocampal gyrus; higher BOLD response to prediction error in the fusiform gyrus, inferior temporal gyrus, and temporal pole; and higher BOLD response to anticipation in the striatum (caudate and putamen), regions near the insula (Rolandic operculum and superior temporal pole), medial frontal cortex (paracentral lobule and orbital part of the superior frontal gyrus), inferior parietal gyrus, postcentral gyrus, anterior cingulate cortex, and superior occipital gyrus.

**Figure 3.**
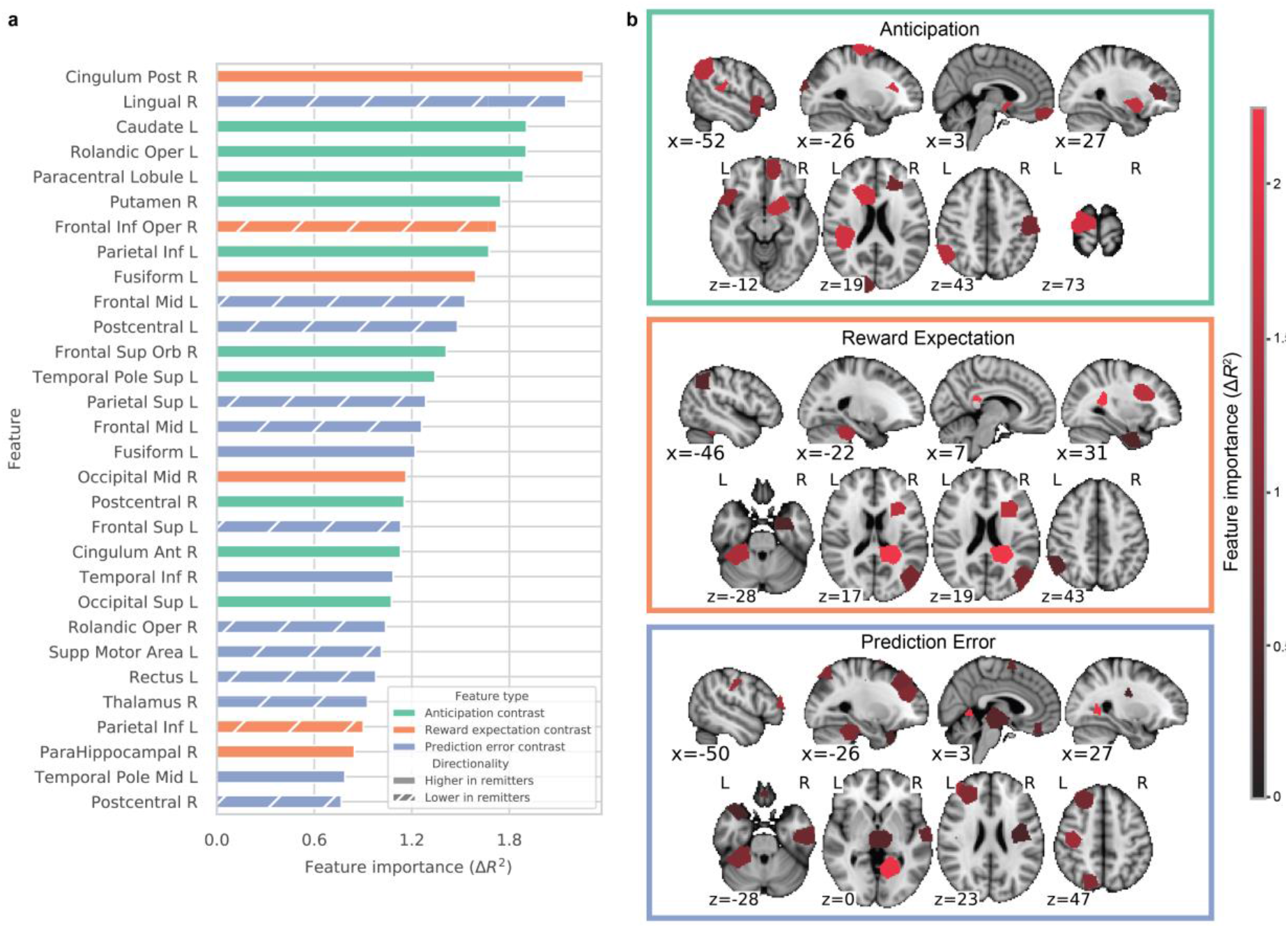
The thirty most important features learned by the bupropion outcome prediction model, as determined by permutation feature importance. **a)** Features are ranked by descending importance and colored by feature type. The directionality of each feature was computed by comparing the mean feature values of subjects who did and did not experience remission; features with lower values in remitters have hatched bars. **b)** The same features are visualized as colored ROIs in the study-specific brain atlas and overlaid on the MNI brain template. Brighter red indicates greater feature importance.

### Treatment Specificity of the Predictive Models

To test whether the models had identified composite predictive biomarkers that are *specific to each treatment*, the model for each treatment was used to predict outcomes for participants from the other two treatment arms. In each case, the predictive performance was low (negative *R*^2^), confirming that each had learned predictive features *specific* to each treatment.

### Comparisons with Traditional Statistical and Deep Learning Methods

Analyses using traditional statistical parametric mapping and classical machine learning approaches found weaker associations and poorer predictive capability (Supplement, Appendix II), reinforcing the need for deep learning. The best performing classical machine learning model was an XGBoost regressor with an *R^2^* of 11% and RMSE of 6.54 for sertraline.

## DISCUSSION

These results describe the first deep learning predictive models for three treatments (sertraline, placebo and bupropion). All 3 models explained a substantial proportion of the variance in ΔHAMD, and NNT for predicting remission ranged from 2.35 for bupropion to 4.31 for sertraline, on par with or lower than the NNTs of many depression treatments^15^. The higher performance of the bupropion model compared to the sertraline model is likely due to the cross-over design of the study; participants were switched to bupropion after failing sertraline in Phase 1, resulting in a smaller and likely less heterogenous cohort. Importantly, remission rates were similar across the 3 treatments at the *group* level: 39% for sertraline, 33% for placebo, and 32% for bupropion. This reinforces the need for a predictive model that can predict *individual* outcomes and preemptively identify the minority of individuals who will respond to an antidepressant.

Higher prediction accuracy was achieved compared to previously published predictors of individual antidepressant treatment outcome. Etkin et al. developed a predictor of escitalopram treatment outcome using cognitive and emotional behavioral measures and achieved an NNT of 3.8, but their results were limited to cognitively-impaired individuals^16^. Gordon et al. analyzed the same data and developed predictors for escitalopram and venlafaxine with NNTs of 2.7 and 4.6, respectively^17^. Gordon et al. also developed a predictor for sertraline with NNT of 3.5, but PPV was 43%, lower than the 62% of the sertraline model developed in this current study. This lower PPV limits clinical utility, as the majority of predicted sertraline responders would be false positives. Of note, no previous studies have developed models to predict bupropion outcome.

### Interactions between sertraline and the reward processing task

The ability of reward processing to predict bupropion outcome is consistent with the primary effect of bupropion on dopamine, the principal neurotransmitter in reward processing. Interestingly, reward processing data were also predictive for sertraline and placebo outcomes. In both cases however, clinical features improved these models, whereas clinical features decreased the accuracy of the bupropion model. This suggests that objective biologic data (e.g. neuroimaging) may act as a better predictor when it corresponds mechanistically to the primary action of the treatment.

Unlike other SSRIs, sertraline has known inhibitory activity for the dopamine transporter, particularly at high dosages^18^. This would likely modulate reward processing circuitry. A second possible explanation is that the reward processing task stimulates the serotonergic emotion circuitry to some degree. Given the shared anatomical regions between the reward and emotion processing circuits, such as the anterior cingulate cortex, medial frontal cortex, and amygdala, there is likely an interaction between the reward processing task and the serotonergic circuits modulated by sertraline^19^.

### Examination of learned composite neuroimaging biomarkers

The important imaging features learned by the models can be compared to existing studies of reward processing neural circuitry, MDD pathophysiology, and antidepressant response biomarkers (**Table 2**). In this section, the features are categorized into those that have a previously known neurobiological basis and those that are novel and could warrant further investigation.

**Table 2.**
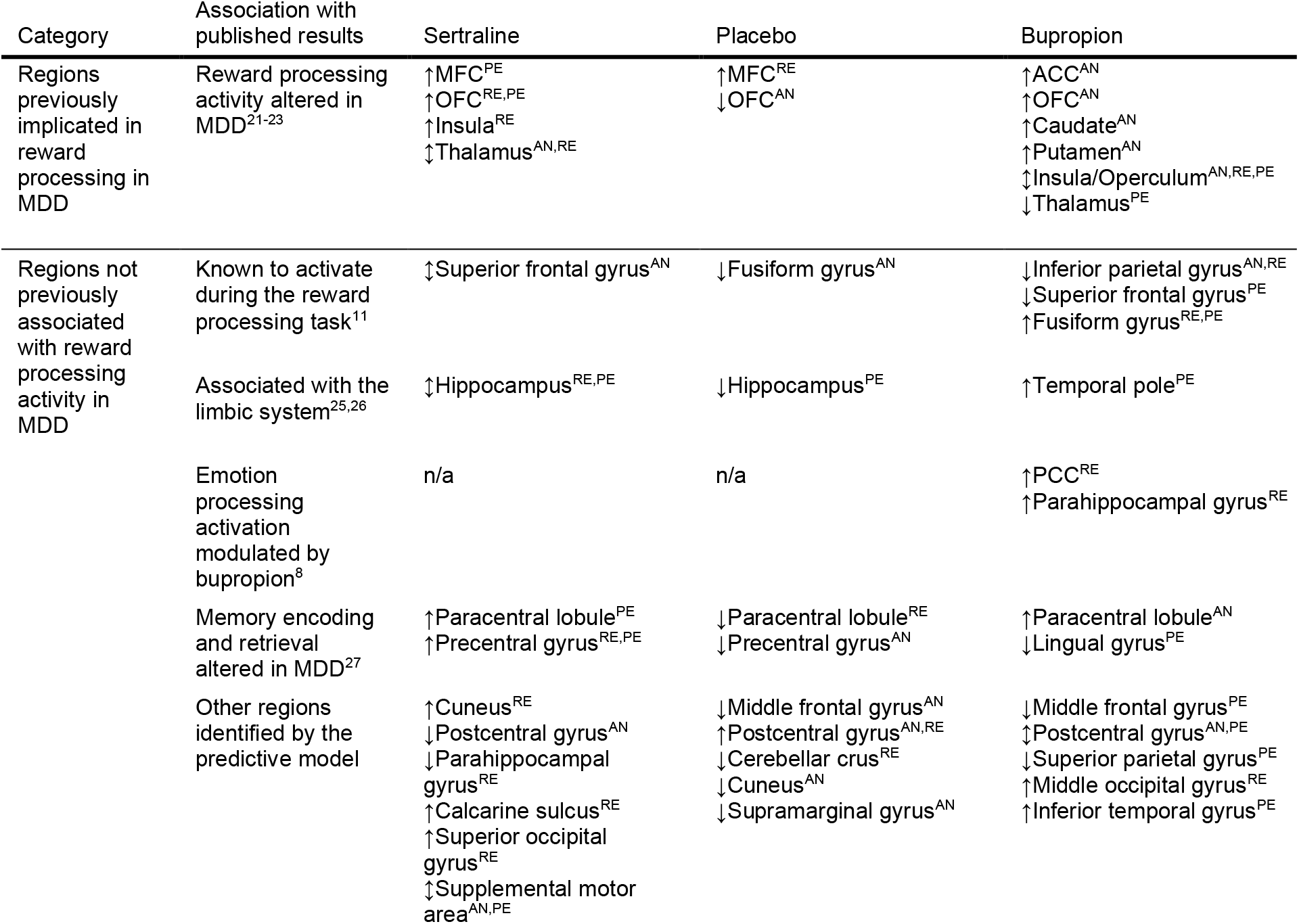
Brain regions identified by the models as containing reward task BOLD activation features predictive of treatment outcome. The regions are separated into two categories. The first category contains regions previously implicated in the context of altered reward processing in MDD or treatment outcome prediction, or known to be activated by the reward processing task used in this study. The second category contains regions that have not been directly associated with reward processing in MDD and that may constitute novel candidate biomarkers of treatment outcome. (↑) indicates that BOLD activation in the region was associated with remission, (↓) indicates association with resistance to treatment, and (↕) indicates that the directionality of the association varied among contrasts. Superscripts indicate which reward task contrast feature was learned by the model in a particular region; *AN*: anticipation, *RE*: reward expectation, *PE*: prediction error. Region abbreviations include the anterior cingulate cortex (ACC), posterior cingulate cortex (PCC), medial frontal cortex (MFC), and orbitofrontal cortex (OFC).

The predictive model for sertraline (**Table 2**, 3^rd^ column) identified regions that have been previously found to have altered BOLD activity during reward processing in MDD, particularly the medial frontal cortex (MFC), orbitofrontal cortex (OFC), insula, and thalamus^19, 20^. While most of these regions were also learned by the bupropion model (**Table 2**, 5^th^ column), the specific predictive contrasts in these regions are distinct between the antidepressants; e.g. prediction error-related activation in the thalamus is predictive of resistance for bupropion while anticipation-related activation in the same region is predictive of remission for sertraline. The placebo model (**Table 2**, 4^th^ column) also learned the MFC and OFC as important regions, but with different predictive contrasts. While the OFC was learned by all three models, higher anticipation-related activation predicted remission for bupropion but resistance for placebo, and higher reward expectation and prediction error-related activation predicted remission for sertraline.

The predictive model for bupropion (**Table 2**, 5^th^ column) also learned several regions that have been previously implicated in altered reward processing in MDD, including the anterior cingulate cortex (ACC), OFC, dorsal striatum (caudate and putamen), insula, and thalamus^20, 19, 21^. Activation in the ACC in response to emotional stimuli has been shown to predict faster improvement in response to fluoxetine, an SSRI^5^, and the current results suggest an additional role in dopaminergic drug response. Also, connectivity between the ACC and the insula was predictive of treatment-independent clinical improvement in an analysis of EEG data from this same study (EMBARC)^22^. Reward task activity in the ventral striatum has been previously identified as a moderator of antidepressant outcome^7^, and a similar moderating relationship is seen here with the dorsal striatum.

Several regions were learned by the models that have not previously associated with altered reward processing in MDD (**Table 2**, bottom). The superior frontal gyrus was learned by both the sertraline and bupropion models, and this region is known to be stimulated by the reward processing task, and consequently would contain a strong signal to be learned by the models^11^. Similarly, the hippocampus, learned by the sertraline and placebo models, and the nearby temporal pole, learned by the bupropion model, are constituents of the limbic system and would likely be activated by reward stimuli^23, 24^. The paracentral lobule was learned by all three models and the precentral gyrus by the sertraline and placebo models. These regions do not have a clear role in the canonical reward processing circuitry but have demonstrated altered activation during memory encoding and retrieval tasks in MDD participants^25^. Additionally, decreased perfusion in the paracentral lobule on arterial spin labelling imaging has been related to antidepressant resistance^26^.

### Synergy between neuroimaging and clinical features

The predictive clinical and demographic features learned by the sertraline and placebo models are described in detail in the Supplement, Appendix III. Of note, higher body mass index predicted resistance to sertraline, while psychomotor agitation predicted remission. For placebo, psychiatric comorbidities, such as anxious distress and panic disorder, and older age predicted resistance. Importantly, *this is one of the first studies to synergize imaging measurements with another modality of information, namely clinical assessments, and this combination yielded stronger predictive signals* in both the sertraline and placebo models. These findings support further investigations into cross-modal composite moderators of treatment response for MDD.

### Limitations

While EMBARC is the largest randomized, placebo-controlled study of antidepressant response with fMRI to date, the enrolled cohort may not fully represent the general population. Inclusion criteria included early onset (before age 30), chronic (episode duration > 2 years), or recurrent (2+ episodes) disease which represents a subpopulation of all MDD patients. In order to most accurately estimate the generalization performance of these predictive models on real-world data, the models should be tested on an additional, independent dataset. In the meantime, the rigorous cross-validation employed here provides high confidence that the results will generalize to additional participants.

Several other studies have found significant biomarkers of antidepressant response with 10-20 participants^27, 5, 8^. However, the 37 participants for bupropion could be considered a limitation of this work. To address this, data augmentation was used to simulate additional fMRI samples. Data augmentation has been shown to enhance the performance of deep learning in data-limited situations^28^. Additionally, the 95% confidence intervals were reported for each *R*^2^ accounting for sample size (*before* data augmentation), indicating that at worst, the model explained 12% of the variance which is still a substantial effect size. Furthermore, permutation testing to check the significance of results also helped to ensure that findings were not spurious.

A secondary limitation for the bupropion treatment group comes from the crossover study design. Participants were selected to receive bupropion after not responding to an initial 8-week course of sertraline treatment, which may have led to a sampling bias for participants less responsive to antidepressants. A follow-up study with a wider treatment design is warranted.

## CONCLUSION

To date, this is the first application of deep learning to prediction of treatment outcome from fMRI in MDD. Deep learning models learned to predict *individual* outcomes for three different treatments and explained a substantial portion of the variance in outcome. Furthermore, these models integrated clinical measurements with imaging data, which has not been extensively explored in other studies of treatment moderators. Important predictive clinical features learned by the models were identified, which can readily inform clinical treatment decision-making. This work also highlights the prognostic value of imaging-based measurements of reward processing. Composite neuroimaging candidate biomarkers were identified for sertraline, bupropion, and placebo, and deeper examination of the important learned features revealed many brain regions in concordance with previous knowledge of MDD neurophysiology as well as several regions not previously implicated in antidepressant research. Future efforts to develop analogous measurements of reward processing in these brain regions, such as through behavioral or other surrogate markers, may form accurate and clinically useful predictive tools for personalized treatment selection. This work is a step towards expediting the selection of appropriate antidepressants, which will help to reduce the need for long and subjective medication trials and to alleviate the considerable morbidity in depression.

## ACKNOWLEDGEMENTS

The EMBARC study was supported by the National Institute of Mental Health of the National Institutes of Health under award numbers U01MH092221 (to M.H.T.) and U01MH092250 (to P.J.M., R.V.P., and M.W.). Valeant Pharmaceuticals donated the Wellbutrin XL used in the study. This work was supported by the EMBARC National Coordinating Center at UT Southwestern Medical Center, with M.H.T. as Coordinating PI, and the Data Center at Columbia and Stony Brook Universities.

## Footnotes

a Structured Clinical Interview for DSM-IV

b Neuroticism-Extraversion-Openness Inventory

c Mood and Anxiety Symptom Questionnaire

d Snaith-Hamilton Pleasure Scale

