## Supplemental Material for "Predictive Patterns of Antidepressant Response from Pre-Treatment Reward Processing using Functional MRI and Deep Learning: Key Results from the EMBARC Randomized Clinical Trial"

**APPENDIX I: SUPPLEMENTAL METHODS**

**Study Design**

Institutional review board approval was obtained from each study site: Columbia University, Massachusetts General Hospital, University of Michigan, and University of Texas Southwestern Medical Center. Written informed consent was obtained from all subjects. Subjects were antidepressant-naïve in the current episode, must have met Structured Clinical Interview for the DSM-IV (SCID) criteria for MDD, and scored ≥ 14 on the Quick Inventory of Depressive Symptomatology (QIDS-SR). Additionally, to reduce heterogeneity, subjects must have had early onset (before age 30), chronic (episode duration > 2 years), or recurrent (2+ episodes) disease. Exclusion criteria included pregnancy, concurrent use of antipsychotics or mood stabilizers, and significant risk of suicide during the study as evaluated by study investigators. Additionally, subjects must not have had a lifetime history of psychosis, bipolar disorder, or epilepsy and must not be receiving depression-specific psychotherapy or somatic treatments.

**Reward Task Paradigm**

The monetary reward task (**Figure S2**) is motivated by differential reactivity to reward anticipation and prediction error, depending on brain region, which has been identified between healthy subjects and MDD subjects. Each trial of the task begins with the *response phase*, during which the subject guesses whether an upcoming number, with possible values of 1-9, will be greater or less than 5. During the *anticipation phase*, the subject is informed about the possible outcome of the current trial. Trials can be "possible win", where the subject wins $1 for a correct guess and loses nothing for a wrong guess, or "possible loss", where the subject loses $0.50 for a wrong guess and wins nothing for a correct guess. During the *outcome phase*, the actual number is revealed, followed by visual feedback indicating whether the subject has won money, lost money, or did not win or lose any money. This is followed by a fixation period before the next trial. A total of 24 trials were conducted, with 12 "possible win" and 12 "possible loss" trials. All subjects received a fixed monetary reward after the task regardless of outcome.

**Data Augmentation**

To improve the performance of the deep learning models and increase their ability to learn the true association between imaging features and treatment outcome, an anatomically-informed data augmentation approach was used to simulate additional fMRI acquisitions. Similar augmentation approaches have demonstrated to dramatically increase the accuracy of deep learning models in natural (non-medical) image applications[1]. In this study, data augmentation was employed to synthesize additional fMRI data from the existing data by simulating variations in brain morphology[2]. Through this approach, each subject's fMRI was coregistered to another randomly selected age- and gender-matched subject's brain to simulate a new fMRI acquisition, with the original subject's fMRI warped onto a new brain morphology. Coregistration target subjects were randomly selected from a different treatment group than the original subject. Because the target subjects are not included in the training, validation, or held-out test data of each treatment-specific model, this avoids biasing the model performance results.

This augmentation increased the effective sample sizes by 500%, providing a total of 222 fMRI images for bupropion, 978 for sertraline, and 696 for placebo. Importantly, this augmented data was used only during model training and not during evaluation. **Table S5** compares the predictive performance achieved with and without data augmentation and demonstrates the benefit of augmentation.

**MRI Preprocessing**

All original and augmented data was preprocessed as follows. Structural MRI were first processed with the ROBEX tool[3] to remove the skull and non-brain voxels. The image is then spatially normalized to the MNI152 T1-weighted template brain using a series of rigid body, affine, and nonlinear symmetric normalization (SyN) registrations in ANTs. This registration method was selected as it has been shown to outperform other registration methods[4]. The normalized sMRI was then segmented into gray matter, white matter, and cerebrospinal fluid with FSL FAST. Functional MRI were corrected for frame-to-frame head motion with FSL MCFLIRT, and frames where the magnitude of head motion was > 1 mm or the Z-score of the intensities was > 3 were marked as outliers to be regressed out during GLM analyses. Brain extraction was performed using the EPI brain extraction method from fMRIPrep, which applies FSL BET and AFNI 3dAutomask and takes the intersection of the two segmentations[5]. Next, spatial normalization was conducted using a direct EPI-based normalization, where the mean functional image frame was directly registered to the MNI152 EPI brain template with ANTs. This direct normalization has been demonstrated to better correct for geometric distortions caused by EPI magnetic inhomogeneities than traditional, T1-based normalization which registers the functional to the structural image and the structural image to the template in two steps[6, 7]. Finally, the normalized fMRI was spatially smoothed with a 6 mm Gaussian filter.

**Contrast Map Computation**

Subject-level generalized linear models (GLMs) were fitted to the fMRI using the SPM12 package. Regressors were defined based on methodology used in prior analyses of this reward task fMRI data[8, 9]. These included regressors for each of the *response, anticipation, outcome,* and *baseline* phases in the task paradigm. Additionally, parametrically modulated regressors were added to represent reward expectation and prediction error. The reward expectation regressor had a value of +0.5 during the *anticipation* phase of “possible win” trials and -0.25 during the *anticipation* phase of “possible loss” trials, which are the expected values of the monetary outcome of these two trial types. The prediction error regressor corresponded to the *outcome* phase and was set to the difference between the outcome and the expected value: +0.5 for a correct guess in a “possible win” trial, -0.5 for a wrong guess in a “possible win” trial, +0.25 for a correct guess in a “possible loss” trial, and -0.25 for a wrong guess in a “possible loss” trial. These 6 primary regressors, their first temporal derivatives, the head motion parameters obtained during preprocessing, and the regressors for the outlier frames were included in the GLM design matrix $\boldsymbol{X}$. White matter and cerebrospinal fluid masks from the sMRI segmentation were applied to mask out unimportant voxels from the analysis. The GLM was fitted:

$$\boldsymbol{Y} = \boldsymbol{X}\boldsymbol{\beta} + \boldsymbol{\epsilon}$$

Where $\boldsymbol{Y}$ is the $\text{time} \times \text{voxels}$ data matrix containing the voxel timeseries, $\boldsymbol{X}$ is the $\text{time} \times \text{regressors}$ design matrix containing the regressor timeseries, $\boldsymbol{\beta}$ is the $\text{regressors} \times \text{voxels}$ matrix containing the fitted coefficients, and $\boldsymbol{\epsilon}$ contains the residuals. The anticipation contrast map was computed as $\beta_{\text{anticipation}} - \beta_{\text{baseline}}$. The reward expectation and prediction error contrast maps were simply $\beta_{\text{reward expectation}}$ and $\beta_{\text{prediction error}}$ respectively.

**Computation of Regional Contrast Values**

Preliminary results showed that a study-specific functional brain atlas, generated from MDD fMRI, yielded superior predictive results when used to extract imaging features from contrast maps compared to a canonical functional brain atlas generated from healthy subjects (Schaefer 2018[10]). A study-specific brain atlas with 200 ROIs was generated from pre-treatment resting-state fMRI images of 283 MDD subjects using the spatially-constrained spectral clustering method developed by Craddock et al[11]. The anatomical label for each ROI was determined by finding the corresponding anatomical structure with the greatest Dice overlap in the widely-used Automated Anatomical Labeling atlas[12]. For each contrast map, including anticipation, reward expectation, and prediction error, the mean of the voxel intensities from the contrast map was computed for each ROI. Concatenation of the 200 mean regional values from each of the 3 contrast maps yielded a vector of 600 fMRI features for each subject.

**Deep Learning Model Training and Hyperparameter Optimization**

To mitigate overfitting, in addition to using the previously described data augmentation, the models were rigorously regularized with L1 and L2 weight regularization, batch normalization, and dropout layers. *Hyperparameters* defining the model architecture, such as number of layers, number of neurons per layer, learning rate, regularization strength, and dropout rate were optimized using a random search. Random search has shown to be an unbiased, highly efficient method for quickly identifying optimal hyperparameters with low computational overhead[13, 14]. With an appropriately defined hyperparameter search space, a random search will find a high-performing model architecture without dependency on the expertise of the investigator. Five hundred hyperparameter configurations were sampled randomly from uniform distributions over predefined hyperparameter ranges (**Table S4**). The same set of 500 candidate models was used for each treatment group. The models were implemented in the Keras and Tensorflow packages and trained using Nvidia Tesla P100 GPUs on the BioHPC computing cluster at UT Southwestern. Models were trained with the Nadam optimizer, with learning rate and decay included as hyperparameters in the random search. The loss function was designed to maximize $R^2$ as was defined as:

$$\ell = \alpha\left(1 - f_{R^2}(y, \hat{y})\right) + \lambda\Omega(\theta)$$

where $f_{R^2}(\cdot)$ computes $R^2$ between the true output values $y$ and the predicted outputs $\hat{y}$ and $\lambda\Omega(\theta)$ is the weight regularization term. The coefficient $\alpha$ was set to 100 empirically to keep the magnitudes of the first and second terms of the loss function within similar ranges.

The predictive performance of each candidate model was validated using nested cross-validation[15]. The data was first split into 3 outer cross-validation folds, stratified by ΔHAMD to ensure representative distributions of subjects in each fold. The training data of each fold was then split again into 5 inner cross-validation folds, which were used to evaluate the performance of each candidate model. For each outer fold, the model with the best $R^2$ across the inner folds was selected, retrained on all inner-fold data of that outer fold, and evaluated on the held-out outer fold data. The mean performance of the best model from each outer fold is reported in the results in the main text.

After completing this random search, plots of performance vs. hyperparameter values were visualized to ensure that local maxima of performance had been observed. This verified that sufficiently large hyperparameter ranges had been searched to identify a high-performing model.

**Computation of Number-Needed-to-Treat**

In this work, the number-needed-to-treat (NNT) is defined as the number of individuals that must be screened by a predictive model to identify one additional remitter or responder, compared to the overall remission or response rate of the treatment in this study:

$$NNT = \frac{1}{r_e - r_c}$$

For example, to compute the NNT for predicting remission, the experimental event rate $r_e$ is the true remission rate in the subjects predicted by the model to remit:

$$r_e = \frac{\text{\# true remitters}}{\text{\# predicted remitters}}$$

And the control event rate $r_c$ is the overall remission rate of the treatment group in the study:

$$r_c = \frac{\text{\# remitters}}{\text{\# subjects in treatment group}}$$

Additionally, a second NNT can be defined as the number of individuals that must be screened to identify one additional remitter or responder, relative to a clinician's performance in making the same treatment selection decisions. The typical antidepressant response rate in clinical practice is estimated to be about 45%[16]. This can be used to define

$$NNT_{clin} = \frac{1}{r_e - r_{clin}}$$

where the control event rate is now

$$r_{clin} = 45\%$$

$NNT_{clin}$ for the 3 predictive models is reported in Appendix II.

**Permutation Testing**

The statistical significance of the model performance results was measured using permutation testing, which tests the null hypothesis that the model did not learn the association between the data and the prediction target[17]. In this approach, a null distribution is generated by permuting the target labels, i.e. ΔHAMD in this study. Specifically, the labels were randomly permuted 500 times and the model was refit and evaluated each time. The *p*-value for each performance metric was obtained by computing the cumulative density function of the null distribution at the actual model performance.

**Feature Importance**

For the best performing models of each random search, important learned features were identified using permutation feature importance[18]. This method leaves the labels undisturbed while each individual feature is permuted among the subjects, ablating any useful information in that feature, and the change in model performance ($R^2$) was measured. This process is then repeated for each feature. Features that incur a greater decrease in performance when permuted are more important for the model's prediction[18]. Within each outer cross-validation fold, features were ranked by importance, and the mean importance rank over all folds was used to identify the most important features learned by the model.

### APPENDIX II: SUPPLEMENTAL RESULTS

**Comparison to Traditional Statistical Analysis and Classical Machine Learning**

A traditional voxel-wise analysis using statistical parametric mapping was performed to identify any group differences in reward-related activation between treatment responders and non-responders. The following group-level comparisons were conducted: responders vs. non-responders, remitters vs. non-remitters, and top quartile of ΔHAMD vs. bottom quartile of

ΔHAMD. None of these comparisons identified significant group differences after false discovery rate correction at $p < 0.05$. These results underscore the importance of using a more statistically powerful analysis such as those described in the preceding sections.

To further evaluate the need for advanced machine learning models, a number of classical machine learning methods were tested to compare the current deep learning approach to other multivariate regression models. The model types tested included multiple linear regression, elastic net regression, K-nearest neighbors, support vector machine, and random forest. Hyperparameter models were optimized and performance on held-out data was evaluated using the same approach that was used on the deep learning models. The same data augmentation was applied and models were tested with imaging features alone and with both imaging and clinical features. The best performance achieved with these classical machine learning models was an $R^2$ of 11% and RMSE of 6.54 in predicting ΔHAMD for sertraline, and results for bupropion and placebo were poor. Compared to the deep learning models, these models were unable to learn to predict treatment outcome from the data with high accuracy.

**Alternate Computation of Number-needed-to-treat**

In the main text, the reported NNT values are computed relative to the actual remission or response rates in each treatment group of the study (see Appendix I, Computation of Number-needed-to-treat). Because treatment assignment was randomized in this study, this NNT may be less relevant to real-world clinical practice. A second metric, $NNT_{clin}$, was computed to compare the performance of these models to clinician performance for the same antidepressant selection decisions. Using an estimated medication response rate of 45% in clinical practice[16], $NNT_{clin}$ was 4.35 for sertraline and 3.85 for bupropion. This indicates that a clinician would need to screen about 4 individuals using the predictive models to identify one additional individual who could be treated successfully with sertraline or bupropion, compared to current clinician decision-making.

### APPENDIX III: SUPPLEMENTAL DISCUSSION

#### Examination of Clinical and Demographic Biomarkers

For the sertraline and placebo models, clinical and demographic features were found to be complementary with imaging features for achieving high predictive performance. Clinical features alone, however, yielded poor predictive performance with $R^2$ of 1.9% for sertraline and 1.8% for placebo (**Table S6**). Additionally, using imaging features alone also provided low predictive performance ($R^2$ < 10%, **Table S5**). For the sertraline model, a small number of clinical measurements were found to be highly important features for predicting treatment outcome (6 out of the top 30 features, **Fig. 1a** in Main Text). Pre-treatment HAMD score was the most important feature overall, with a higher total score on the either the 17-item or 24-item version predicting treatment resistance. Higher body mass index (BMI) also predicted resistance, which is corroborated by other studies showing that obesity correlates with poorer outcomes with SSRIs[19, 20]. A notable finding was that concurrent psychomotor agitation predicted remission. Sertraline is known to effectively treat psychomotor agitation, compared to other SSRIs such as fluoxetine, and a prior study saw a non-significantly higher response rate to sertraline vs. nortriptyline in agitated subjects[21, 22].

Regarding the placebo model, several associated symptoms and psychiatric comorbidities were important features predictive of resistance, such as concurrent anxious distress, concurrent panic disorder, and anhedonia. As with sertraline, pre-treatment comorbidity score was a highly important feature for predicting resistance. Temporal features also appeared to be predictive, including older age at time of study being predictive of resistance and a longer period of time without dysphoria being predictive of remission. Asian and Caucasian race were both learned as predictive features, with Asian subjects being more likely to remit, but this may be an artifactual finding given that only 7% of the placebo treatment group (8 subjects) was Asian.

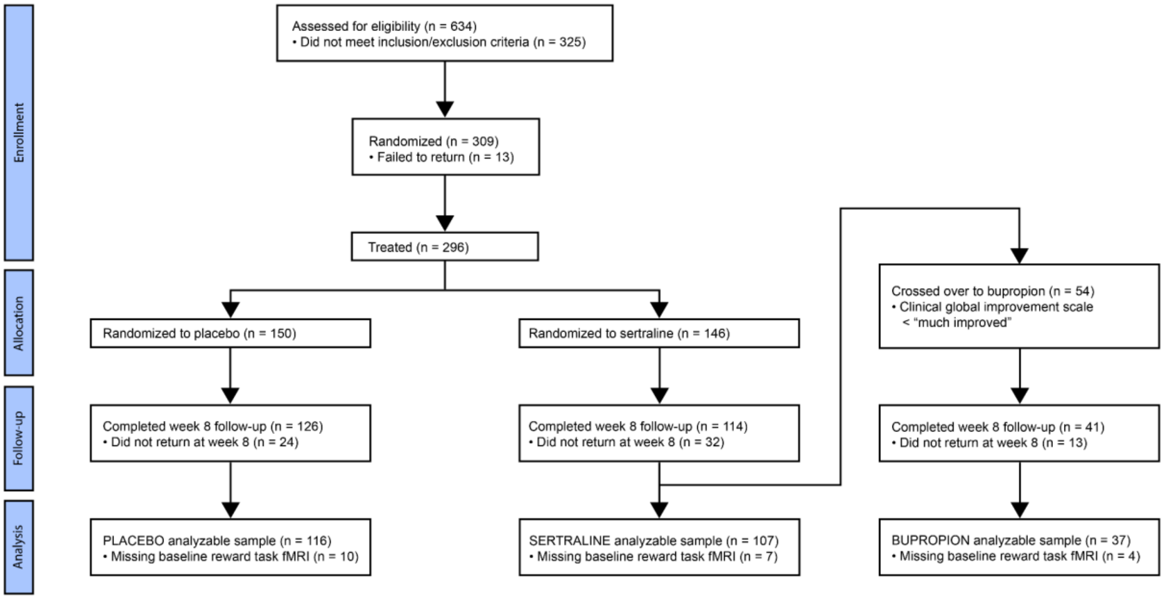

**Figure S1**. CONSORT diagram of study participants.

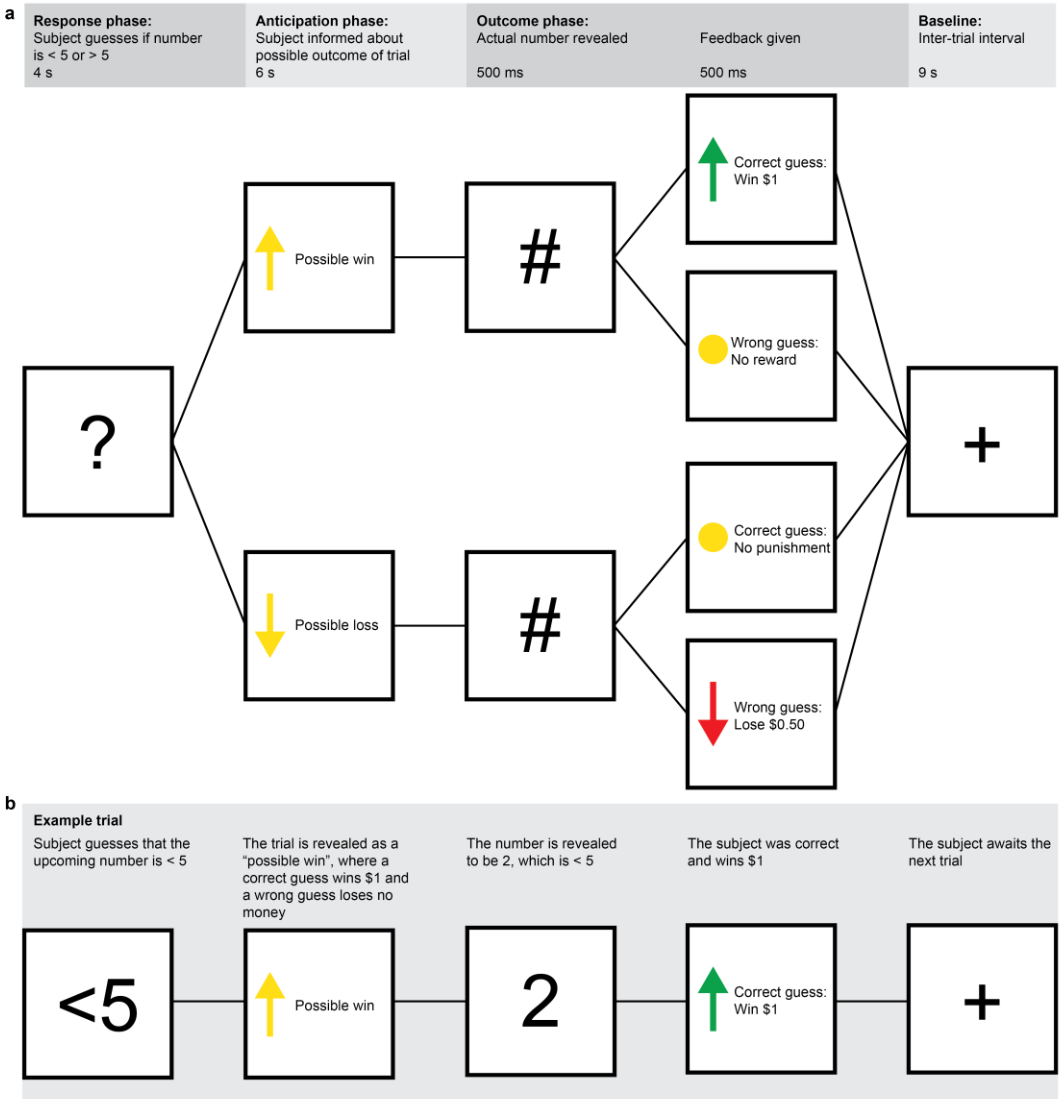

**Figure S2**. Block-design reward task paradigm employed in this study. The task lasts 8 minutes and includes 24 trials. **a**) Flowchart demonstrating the possible stimuli and outcomes for a single trial. In each trial, the subject guesses whether the upcoming number (1-9) is greater or less than 5. They are shown whether the trial is a "possible win" with a reward for a correct guess or a "possible loss" with a punishment for a wrong guess, and the outcome is then presented. **b**) Diagram for an example trial. In this case, the subject guesses that the upcoming number is less than 5, and the trial is a "possible win". The actual number is 2, and the subject receives $1 for a correct guess.

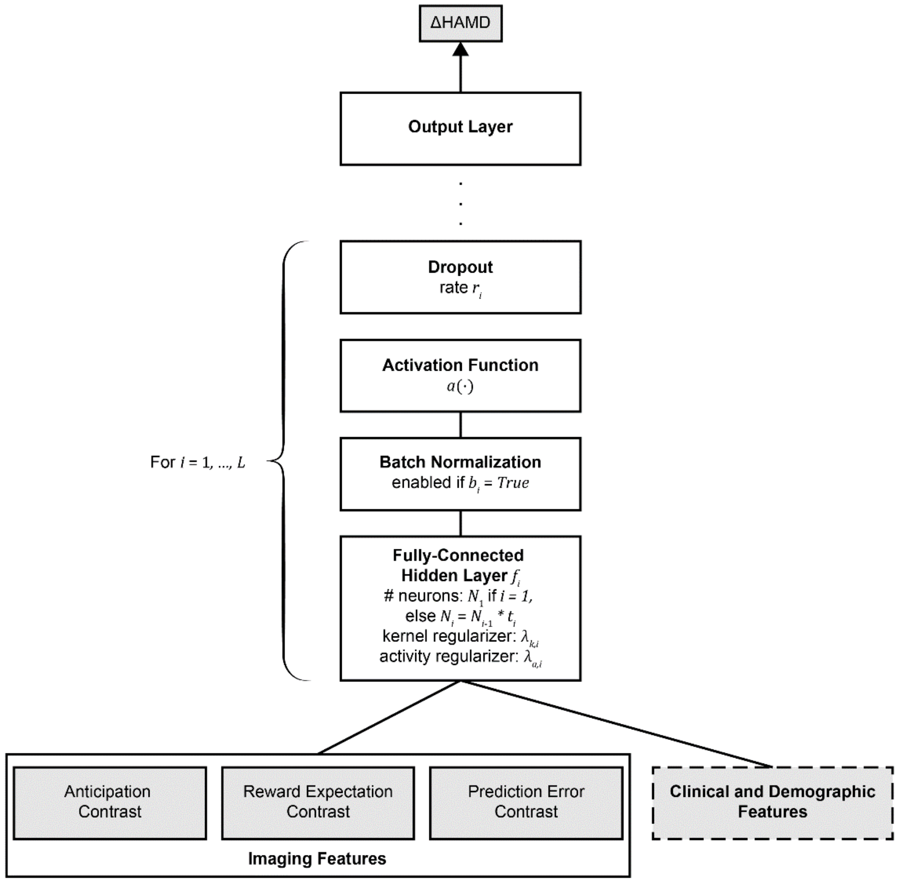

**Figure S3**. Schematic for the feed-forward neural networks developed in this work. Hyperparameters are indicated for each layer and were optimized using a random search for each treatment. Ranges of hyperparameters that were searched are listed in **Table S3**. Inputs to these models included imaging features, extracted from the contrast maps for each of the three task conditions, and clinical and demographic features for the sertraline and placebo models. This data is fed through a series of fully-connected hidden layers $f_i$ for $i = 1, \ldots, L$, and the number of layers $L$ was optimized during the random search. Regularization parameters $\lambda_{k,i}$ and $\lambda_{a,i}$, use of batch normalization $b_i$, and dropout rate $r_i$ for each layer were included as optimized hyperparameters. The activation function $a(\cdot)$ for all layers was an additional hyperparameter. The final output layer returns the ΔHAMD prediction.

**Table S1.** Demographic, pre-treatment clinical characteristics, and 8-week treatment outcomes for sertraline, placebo, and bupropion treatment groups.

| | Sertraline | | Placebo | | Bupropion | |
|---|---|---|---|---|---|---|
| **Total subjects** | 106 | | 116 | | 37 | |
| **Demographics** | | | | | | |
| Female | 73 | 69% | 73 | 63% | 26 | 70% |
| Race | | | | | | |
| White | 72 | 68% | 83 | 72% | 24 | 65% |
| African American | 20 | 19% | 17 | 15% | 8 | 22% |
| Asian | 5 | 5% | 8 | 7% | 3 | 8% |
| Other | 9 | 8% | 8 | 6% | 2 | 5% |
| Hispanic | 19 | 18% | 22 | 19% | 6 | 16% |
| Employed | 61 | 58% | 69 | 59% | 20 | 54% |
| Age | 38.38 ± 13.95 | | 37.40 ± 12.80 | | 37.51 ± 14.32 | |
| **Clinical characteristics** | | | | | | |
| Age of first major depressive episode | 16.16 ± 5.86 | | 16.57 ± 5.91 | | 16.11 ± 5.77 | |
| Pre-treatment HAMD | 18.60 ± 4.45 | | 18.57 ± 4.26 | | 18.00 ± 3.96 | |
| **Treatment outcomes** | | | | | | |
| ΔHAMD (Week 8 – pre-treatment) | 7.89 ± 7.16 | | 6.70 ± 6.93 | | 5.46 ± 5.57 | |
| Remission | 41 | 39% | 38 | 33% | 12 | 32% |
| Response | 57 | 54% | 41 | 35% | 15 | 41% |

**Table S2**. Scanner and pulse sequence information for each study site.

| | **Columbia University** | **Massachusetts General Hospital** | **University of Michigan** | **UT Southwestern Medical Center** |
|---|---|---|---|---|
| Scanner | General Electric Signa HDx 3T | Siemens TrioTim 3T | Philips Achieva 3T | Philips Ingenia 3T |
| Structural MRI | | | | |
| Sequence | FSPGR | MPRAGE | TFE | MPRAGE |
| TR/TI/TE | 6.0ms/900ms/2.4ms | 2300ms/900ms/2.54ms | 8.2ms/1100ms/3.7 ms | 2100ms/1100ms/3.7ms |
| Flip angle | 9° | 9° | 12° | 12° |
| Dimensions | 256 x 256 x 174 | 256 x 256 x 176 | 256 x 256 x 178 | 256 x 256 x 178 |
| Voxel size | 1 x 1 x 1 mm | 1 x 1 x 1 mm | 1 x 1 x 1 mm | 1 x 1 x 1 mm |
| Functional MRI | | | | |
| Sequence | GE-EPI | GE-EPI | GE-EPI | GE-EPI |
| TR/TE | 2000ms/28ms | 2000ms/28ms | 2000ms/28ms | 2000ms/28ms |
| Flip angle | 90° | 90° | 90° | 90° |
| Dimensions | 64 x 64 x 39 | 64 x 64 x 39 | 64 x 64 x 39 | 64 x 64 x 39 |
| Voxel size | 3.2 x 3.2 x 3.1 mm | 3.2 x 3.2 x 3.1 mm | 3.2 x 3.2 x 3.1 mm | 3.2 x 3.2 x 3.1 mm |
| Dummy scans | 5 | 5 | 5 | 5 |
| Number of volumes, reward task | 240 | 240 | 240 | 240 |
| Total acquisition time, resting state | 480 s | 480 s | 480 s | 480 s |
| Number of volumes, resting state | 180 | 180 | 180 | 180 |
| Total acquisition time, resting state | 360 s | 360 s | 360 s | 360 s |

**Table S3.** Clinical features used as inputs for sertraline and placebo predictive models.

| Clinical assessment name | Items used |
|---|---|
| Body mass index | |
| Clinical history | Number of suicide attempts, lifetime suicide rating |
| 17-item Hamilton Rating Scale for Depression ($HAMD_{17}$) | Total |
| 24-item Hamilton Rating Scale for Depression ($HAMD_{24}$) | Total |
| Altman Self-Rating Mania Scale (ASRM) | Total |
| Anger Attack Questionaire (AAQ) | Total |
| Childhood Trauma Questionnaire (CTQ) | Emotional Abuse, Emotional Neglect, Physical Abuse, Physical Neglect, Sexual Abuse, and Validity subscores |
| Columbia Suicide Severity Rating (CSSRS) | Baseline intensity score |
| Concise Health Risk Tracking (CHRTP) | Propensity score, risk score |
| Edinburgh Handedness Inventory (EHI) | Total |
| Fagerstrom Test of Nicotine Dependence (FTND) | Current cigarette-smoking status |
| Family History Screen (FHS) | All items |
| Mood and Anxiety Symptoms Questionnaire (MASQ) | Anxious Arousal, Anhedonic Depression, and General Distress subscores |
| Mood Disorders Questionnaire (MDQ) | Total |
| NEO-Five Factor Inventory | Agreeableness, Conscientiousness, Extraversion, Neuroticism, and Openness subscores |
| 16-item Quick Inventory of Depressive Symptomatology (QIDS SR16) | Total |
| Structured Clinical Interview for DSM-5 (SCID) | Current episode duration<br>Current episode specifier (melancholic, atypical, or catatonic)<br>Number of episodes<br>Presence of anxious distress, mixed features, insomnia, hypersomnia, psychomotor agitation, psychomotor retardation<br>History of alcoholism, generalized anxiety, bipolar disorder, panic disorder, or psychotic symptoms |
| Self-Administered Comorbidity Questionnaire (SCQ) | Total |
| Snaith-Hamilton Pleasure Scale (SHAPS) | Ordinal and dichotomous total |
| Social Adjustment Scale (SAS) short form | Total and mean |
| Speilberger State Anxiety Inventory (STAI) | Pre-fMRI and post-fMRI score |
| Standard Assessment of Personality Abbreviated Scale (SAPAS) | Total |
| Visual Analog Mood Scales (VAMS) | Happy-sad, quick witted, relaxed-tense scores |

**Table S4.** Hyperparameter ranges used during hyperparameter optimization. Hyperparameter values were selected uniform randomly from these ranges to create 500 model configurations, which were tested with nested cross-validation to identify an optimal model configuration for the predictive task for each treatment. A model schematic is illustrated in **Figure S1** and is labelled accordingly with these hyperparameters.

| **Hyperparameter** | **Possible values** |
|---|---|
| Layer hyperparameters | |
| Number of fully-connected hidden layers, $L$ | 1, 2, 3 |
| Number of neurons in the first hidden layer, $N_1$ | 64, 96, 128, …, 512 |
| Kernel regularizer, $\lambda_{k,i}$ | L1, L2, L1+L2 |
| Activity regularizer, $\lambda_{a,i}$ | L1, L2, L1+L2 |
| % decrease in hidden layer size from previous layer, $t_i$ | 50%, 75% |
| Batch normalization, $b_i$ | True, False |
| Dropout rate, $r_i$ | 0.3, 0.4, 0.5, …, 0.9 |
| Activation function, $a(\cdot)$ | ReLU, LeakyReLU, ELU, PReLU |
| Nadam Optimizer hyperparameters | |
| Learning rate | 0.001, 0.0011, 0.0012, …, 0.003 |
| Schedule decay | 0.003, 0.0035, 0.004, …, 0.006 |

**Table S5**. Treatment outcome prediction performance for each treatment group, with and without clinical demographic features and with or without data augmentation. Rows with **bold** text are the results presented in the main text. Performance metrics are coefficient of determination ($R^2$) and root mean squared error (RMSE) for predicting the numerical target of ΔHAMD. For predicting the binary targets of remission and response, performance metrics include number-needed-to-treat (NNT), positive predictive value (PPV), and area under the receiver operating characteristic curve (AUROC). NNT of N/A indicates that the model did not achieve better than chance accuracy for that particular prediction target.

| | | | Prediction target | | | | | | | |
|---|---|---|---|---|---|---|---|---|---|---|
| | **Features** | **Augmentation** | **ΔHAMD** | | **Remission** | | | **Response** | | |
| **Treatment** | **used** | **used** | $R^2$ | RMSE | NNT | PPV | AUROC | NNT | PPV | AUROC |
| Sertraline | Imaging | No | 2% | 6.91 | 11.61 | 0.51 | 0.56 | N/A | 0.48 | 0.49 |
| | Imaging | Yes | 1% | 6.96 | 7.56 | 0.56 | 0.57 | N/A | 0.42 | 0.46 |
| | Imaging, clinical | No | 10% | 6.59 | 5.66 | 0.60 | 0.61 | 13.42 | 0.57 | 0.55 |
| | **Imaging, clinical** | **Yes** | **35%** | **5.75** | **4.31** | **0.62** | **0.60** | **4.88** | **0.68** | **0.63** |
| Placebo | Imaging | No | 1% | 6.86 | 3.84 | 0.59 | 0.59 | N/A | 0.35 | 0.50 |
| | Imaging | Yes | 7% | 6.65 | 7.47 | 0.46 | 0.53 | 36.36 | 0.38 | 0.51 |
| | Imaging, clinical | No | 8% | 6.60 | 4.26 | 0.56 | 0.57 | 14.79 | 0.42 | 0.52 |
| | **Imaging, clinical** | **Yes** | **23%** | **6.06** | **2.78** | **0.69** | **0.61** | **3.19** | **0.67** | **0.64** |
| Bupropion | Imaging | No | 30% | 4.60 | 3.63 | 0.60 | 0.67 | 5.14 | 0.60 | 0.55 |
| | **Imaging** | **Yes** | **37%** | **4.36** | **2.35** | **0.75** | **0.71** | **3.24** | **0.71** | **0.62** |
| | Imaging, clinical | No | 25% | 4.74 | 2.66 | 0.70 | 0.73 | 43.17 | 0.43 | 0.51 |
| | Imaging, clinical | Yes | 14% | 5.09 | 2.61 | 0.63 | 0.65 | N/A | 0.38 | 0.49 |

**Table S6**. Treatment outcome prediction performance using clinical features only and traditional machine learning models. Lasso regression, ridge regression, elastic net regression, support vector machine (with linear, radial basis function, and polynomial kernels), random forest, AdaBoost, and Gradient Boosting were all tested for this predictive task. Hyperparameters of each model type were optimized using random search with 100 tested hyperparameter configurations. Generalization performance was measured using nested cross-validation with 10 inner and 10 outer folds. Across all the models, the highest predictive performance observed was $R^2$ of 1.9% in predicting sertraline outcome.

| | | **ΔHAMD prediction performance** | |
|---|---|---|---|
| **Treatment** | **Model type** | $R^2$ | RMSE |
| Sertraline | Lasso regression | 1.9% | 6.93 |
| Placebo | Ridge regression | 1.8% | 6.93 |
| Bupropion | SVM, radial basis function kernel | < 0 | 6.16 |
